# Integrated Stress Response and Necroptosis Drive Epithelial Dysfunction in Crohn’s Disease: Repurposing Cancer drugs for Permeability Barrier Healing

**DOI:** 10.1101/2025.09.16.676680

**Authors:** Debasish Halder, Ritwika Biswas, Amin Esmaeilnia, Jason Ken Hou, Yaohong Wang, Seema Khurana

## Abstract

**Background and Aims:** Epithelial permeability barrier dysfunction is a central pathogenic driver of Crohn’s disease (CD), fueling microbial translocation, chronic inflammation, and progressive tissue injury. While current therapies suppress inflammation, none directly restore epithelial barrier function. Importantly, in CD patients, permeability barrier healing (BH) rather than mucosal healing is associated with long-term remission and a reduced risk of disease complications. Yet BH remains an unaddressed therapeutic target in CD. Here, we investigated whether pharmacologic inhibition of the integrated stress response (ISR) and RIPK3-mediated necroptosis, two convergent pathways of epithelial injury, can promote epithelial viability, regeneration, and barrier integrity in CD.

**Methods:** We employed villin-1/gelsolin double knockout (DKO) mice with epithelial-intrinsic ISR activation, TNF^ΔARE/+^ mice with chronic inflammation, and CD patient-derived enteroids (PDEs). Animals and PDE were treated with ISR inhibitor ISRIB, RIPK3 inhibitor Necrostatin-1 (Nec-1), or FDA-approved cancer drugs pazopanib and ponatinib, repurposed as potent RIPK3 inhibitors. Epithelial survival, regenerative growth (enteroid formation, budding), and barrier function (transepithelial electrical resistance, TEER) were assessed.

**Results:** Chronic ISR activation and necroptosis were prominent in both murine models and CD PDEs, causing epithelial death, Paneth cell expansion, impaired enteroid survival, and regenerative failure. Pharmacologic inhibition with ISRIB, Nec-1, pazopanib, or ponatinib restored villus architecture, reduced inflammation, enhanced epithelial survival and regeneration, and significantly improved TEER.

**Conclusions:** ISR activation and RIPK3-mediated necroptosis converge to drive epithelial injury and barrier dysfunction in CD. Repurposing pazopanib and ponatinib restored epithelial regeneration and BH, offering an immediately translatable therapeutic strategy for sustained remission in CD.

**Synopsis:** ISR activation and RIPK3-mediated necroptosis drive epithelial injury in Crohn’s disease. Repurposed RIPK3 inhibitors, pazopanib and ponatinib, restore epithelial homeostasis and permeability barrier function, providing a translational strategy to achieve sustained remission in CD.

## Introduction

Crohn’s disease (CD) is a chronic, relapsing inflammatory disorder of the gastrointestinal tract characterized by transmural inflammation, progressive mucosal damage, and the development of complications such as strictures, fistulae, and abscesses. Despite major therapeutic advances with biologics and small molecules, most current CD therapies primarily target immune pathways and not re-epithelialization of the damaged gut. Although inflammation control improves symptoms it achieves mucosal healing or more appropriately endoscopic remission (ER), defined as the absence of inflammation, ulcers, or bleeding in the gastrointestinal tract, in a modest 15-20% of CD patients.[1] While endoscopic remission is an important treatment goal in CD associated with improved clinical outcomes, nonetheless a vast majority of CD patients remain vulnerable to disease progression and intestinal complications.[1] More notably, the presence of mucosal healing in CD does not always correspond with clinical remission, or the resolution of a patient’s symptoms.[2] Additionally, a proportional clinical benefit is not received by most CD and are instead exposed to the risks of potent immunosuppressive therapies.

A defining feature of CD pathogenesis is intestinal permeability barrier dysfunction.[3] Under homeostatic conditions, the intestinal epithelium provides a dynamic and selective permeability barrier that separates luminal microbes from the mucosal immune system. Loss of barrier integrity permits the translocation of commensal and pathogenic bacteria as well as antigens or other foreign substances from the gut lumen into the underlying tissue, triggering an exaggerated and uncontrolled immune response.[4] In CD, this sets in motion a vicious cycle where microbial translocation drives inflammation, inflammation induces further epithelial injury, and the subsequent chronic tissue damage leads to fibrostenosis and/or fistulization, which ultimately increases the need for hospitalization and surgery.[5] Epidemiologic and prospective studies have demonstrated that permeability barrier dysfunction is not merely a consequence of inflammation but an initiating factor in CD.[3, 6] Specifically, asymptomatic individuals with increased intestinal permeability are at significantly higher risk of developing CD, providing causal evidence that permeability barrier dysfunction contributes to disease initiation.[6] Relatives of CD patients often display increased intestinal permeability even before clinical disease onset, supporting a heritable component of permeability barrier defects as well as identify barrier dysfunction as a primary defect in CD.[7, 8] Together these findings highlight the etiologic and therapeutic importance of epithelial permeability barrier restoration in CD.[5, 9]

Therapeutic goals in CD have historically been framed around symptom control, corticosteroid-free remission, and mucosal healing.[10] Indeed, the STRIDE-II consensus reinforced “deep remission,” defined as clinical remission plus ER, as the gold standard treatment target in CD.[11] Achieving ER in CD has been correlated with reduced hospitalization and lower surgical risk.[10, 12] However, emerging evidence shows that compared to ER, permeability barrier healing (BH) is a superior predictor of sustained remission and more favorable long-term outcomes in CD.[13, 14] The prospective ERICa trial demonstrated that BH, outperformed ER and histologic remission in predicting adverse outcomes such as intestinal complications, hospitalization and need for surgery.[2] BH therefore represents a more meaningful therapeutic endpoint in CD. While ER reflects inflammation control and mucosal healing, BH captures restoration of the fundamental epithelial permeability barrier defect that drives CD pathology.[3] Despite this, no approved CD therapy directly targets epithelial permeability barrier repair. This therapeutic gap underscores the urgent need for permeability barrier-focused interventions that complement existing immunosuppressive strategies in CD.

The integrated stress response (ISR) is a highly conserved adaptive pathway that regulates cellular responses to diverse stressors including nutrient deprivation, endoplasmic reticulum stress, mitochondrial dysfunction, microbial and viral infections.[15] Central to ISR signaling is phosphorylation of eukaryotic initiation factor 2α (peIF2α), which transiently attenuates global protein translation while allowing selective translation of stress-adaptive genes.[15] Acute ISR activation is protective, conserves resources and promotes cell survival.[15] In contrast, chronic or unresolved ISR signaling promotes autophagy, or programmed cell death *via* apoptosis or necroptosis, shifting the balance from adaptation to injury.[15, 16] Persistent ISR activation is implicated in the etiology of several neurodegenerative disorders including Alzheimer’s and Parkinson’s disease and manipulation of ISR is emerging as a promising therapeutic approach.[17] Resolution of ISR requires dephosphorylation of peIF2α by GADD34-PP1c phosphatase complex, a process sensitive to cellular actin dynamics.[18] Prior work from our group has shown that villin-1 and gelsolin, two intestinal epithelial cell (IEC) actin-severing proteins, regulate PP1c activity.[16] Their loss leads to sustained ISR activation, mitochondrial stress, and IEC death *via* necroptosis.[16] Double knockout (DKO) mice lacking villin-1 and gelsolin develop spontaneous ileocolitis recapitulating human CD pathology due to chronic ISR signaling, induction of the autophagy protein IRGM1, and IRGM1-induced IEC death by necroptosis.[16] Notably, similar molecular defects are seen in CD patients, including reduced villin-1/gelsolin expression, chronic ISR activation, and IRGM1-induced IEC death by necroptosis, underscoring the relevance of this pathway to the pathophysiology of CD.[16] These findings suggest that therapeutic targeting of the ISR could restore epithelial homeostasis in CD.

Necroptosis is a pro-inflammatory form of programmed cell death that is mediated by receptor-interacting protein kinases RIPK1 and RIPK3 and executed by mixed lineage kinase domain like (MLKL).[19, 20] Beyond depleting IECs, necroptosis leads to the release of damage-associated molecular patterns (DAMPs) which augment intestinal inflammation and epithelial injury.[16] Moreover, independent of its role in necroptosis, RIPK3 is a key regulator of inflammation regulating the activation of NF-kB and the NLRP3 inflammasome.[21, 22] RIPK3 and MLKL levels are increased in inflamed tissues from IBD patients.[23] Additionally, we have shown that CD patients display not only and upregulation of RIPK3 expression, but also higher levels of RIPK3-mediated necroptosis, and increased release of nuclear High Mobility Group Box 1 (HMGB1) protein.[16] The dual roles of RIPK3 in necroptosis and inflammation suggest that pharmacologic inhibition of RIPK3 could attenuate IEC death and pro-inflammatory signaling in CD.

TNF-α is a major contributor to intestinal inflammation that is linked to the pathogenesis of CD.[24] Moreover, genetic studies have identified TNF-α as a CD susceptibility locus while anti-TNF therapy is a mainstay of CD treatment.[24, 25] ISR activation modulates pro-inflammatory cytokine production however it is not known if pro-inflammatory cytokines like TNF-α can act as stressors to activate ISR signaling. In this study, we used two complementary murine models of CD, the villin-1/gelsolin DKO mice, which model epithelial-intrinsic ISR dysregulation, and TNF^ΔARE/+^ mice, which develop chronic inflammation due to TNF-α dysregulation, together with CD patient-derived enteroids (PDEs) to test our hypothesis that ISR activation in response to intrinsic (villin-1/gelsolin loss) or extrinsic (TNF-α) stressors and downstream of it necroptosis, are the main drivers of epithelial dysfunction in CD. Furthermore, using these *in vivo* and *ex vivo* murine and human models of CD, we evaluate our hypothesis that pharmacologic inhibition of ISR and necroptosis can restore epithelial homeostasis and permeability barrier integrity. Our findings establish a mechanistic framework linking ISR and necroptosis to permeability barrier failure in CD and demonstrate the translational potential of repurposed RIPK3 inhibitors as novel therapies to achieve mucosal and permeability barrier healing in CD.

## Methods

### Mouse Models of Crohn’s Disease

The Institutional Animal Care and Use Committee (IACUC) of the University of Houston approved all animal experiments. Double knockout (DKO) mice lacking villin-1 and gelsolin were generated as previously described.[26] TNF^ΔARE/+^ mice, which develop chronic ileitis due to dysregulated TNF-α expression, were obtained from Dr. Fabio Cominelli (Case Western Reserve University).[27] Wild-type (WT) littermates of DKO and the TNF^+/+^ served as controls. Both male and female mice were studied at 20 and 35 weeks of age, corresponding to peak spontaneous ileitis in TNF^ΔARE/+^ and DKO mice, respectively.[16, 28]

### Intestinal Mouse and Human Enteroid Culture

Human CD and non-CD control samples were obtained under protocols approved by the Institutional Review Boards of Baylor College of Medicine and University of Houston. Crypts from murine and human distal ileum were isolated by incubation in 2mM EDTA and enteroids were derived as described before.[29] The organoid forming capacity of stem cell or the organoid forming efficiency (percentage) is defined as a percentage of the number of organoids per number of plated stem cells at day 7, when one cell is added per well.[30] The percentage of budded enteroids was calculated as: (number of enteroids with ≥1 crypt-like bud ÷ total number of enteroids counted) x 100 on day 7.[31] Crypt buds per enteroid, a measure of stem cell-driven proliferation and regeneration, was also quantified as described before.[31] At least 50 enteroids per sample were assessed for all quantitative assays. Epithelial cell mortality was assessed using Hoechst 33342 or propidium iodide staining. Imaging was performed on a Fluoview FV2000 laser scanning confocal microscope with a 60X NA 1.35 objective.

### *In vivo* and *Ex vivo* Treatments

Mice received intraperitoneal injections of Necrostatin-1 (Nec-1; 1.65 mg/kg) or ISRIB (0.25 mg/kg) every other day starting at 1 month of age.[32] Following treatment, TNF^ΔARE/+^ mice were euthanized at 6 months of age and DKO at 8 months of age and ileal tissue collected for histology, immunostaining, and protein analysis. Vehicle-treated animals (0.3% DMSO) served as controls. *Ex vivo*, DKO, TNF^ΔARE/+^ mouse enteroids and CD patient-derived enteroids (PDEs) were treated with ISRIB (50 nM, Cayman Chemicals), Necrostatin-1 (10 μM), pazopanib (0.5 μM), or ponatinib (2 nM) for 7 days and compared with vehicle (DMSO) treated controls.

### Histology and Immunostaining

Distal ileal specimens were fixed in 4% paraformaldehyde and embedded in paraffin. Five-micrometer sections were stained with hematoxylin and eosin (H&E) to evaluate villus architecture, crypt morphology, and inflammatory infiltration. Histological evaluation of intestinal inflammation was scored as described before.[33] For immunohistochemistry, sections were deparaffinized, antigen-retrieved, and incubated with antibodies to phospho-eIF2α, IRGM1, or lysozyme. Images were captured with an Olympus BX53 microscope. For immunoblotting, protein lysates from distal ileum were prepared using RIPA buffer with proteases and phosphatase inhibitors. Equal protein amounts were separated on SDS-PAGE, transferred to PVDF membranes, and probed with antibodies against eIF2α, peIF2α, IRGM1, and β-actin. Bands were visualized using chemiluminescence (Thermo Fisher).

### Transepithelial Electrical Resistance (TEER)

CD PDEs were seeded on Transwell inserts (0.4 μm pore size, Corning) coated with 20 μg/ml collagen type 1 (rat tail) in 0.01% acetic acid. Enteroids were dissociated to single cells using TrypIE^TM^ (Gibco, Thermo Fisher Scientific) for 10 min at 37°C and cultured as monolayers until confluent. TEER was measured using an EVOM2 epithelial voltohmmeter (World Precision Instruments). Each Insert was measured three times; average values were corrected for background and normalized to surface area to calculate net resistance (Ω.cm^2^).

### Statistical Analysis

All experiments were performed with at least three biological replicates. Data are presented as mean ± standard error of mean (SEM). Statistical analyses were conducted using GraphPad Prism v9. Comparisons between two groups were performed using unpaired two-tailed Student’s t-test; comparison of > 2 groups used one-way ANOVA with Tukey’s post hoc test. A p-value of <0.05 was considered statistically significant.

## Results

### ISR activation is a convergent hallmark of epithelial injury in CD models

To determine whether intrinsic epithelial stress or chronic inflammation (extrinsic stressor) converge on epithelial dysfunction and ISR activation, we examined villin-1/gelsolin double knockout (DKO) mice, which model epithelial-intrinsic ISR dysregulation, and TNF^ΔARE/+^ mice, which develop chronic ileitis due to TNF dysregulation.[16, 27] Both strains developed spontaneous ileitis and transmural inflammation resembling CD, as shown before.[16, 27] Histomorphological analysis of distal ileum revealed profound epithelial damage in DKO and TNF^ΔARE/+^ mice, including villus blunting, villus fusion, and crypt elongation, accompanied by thickening of the muscularis propria [Fig 1A-B]. Inflammatory cell infiltration was significantly elevated, as quantified by histological scoring (p<0.0001, n=6 mice per group). Paneth cell hyperplasia and displacement was observed in both mouse models, mirroring phenotypes characteristic of human CD [Fig 1A-B; p<0.0001, n=6 mice per group.].[34] These findings are consistent with expansion of the proliferative compartment, a change in IEC lineage, structural compromise of epithelial integrity, and chronic inflammatory remodeling.

**Figure 1.**
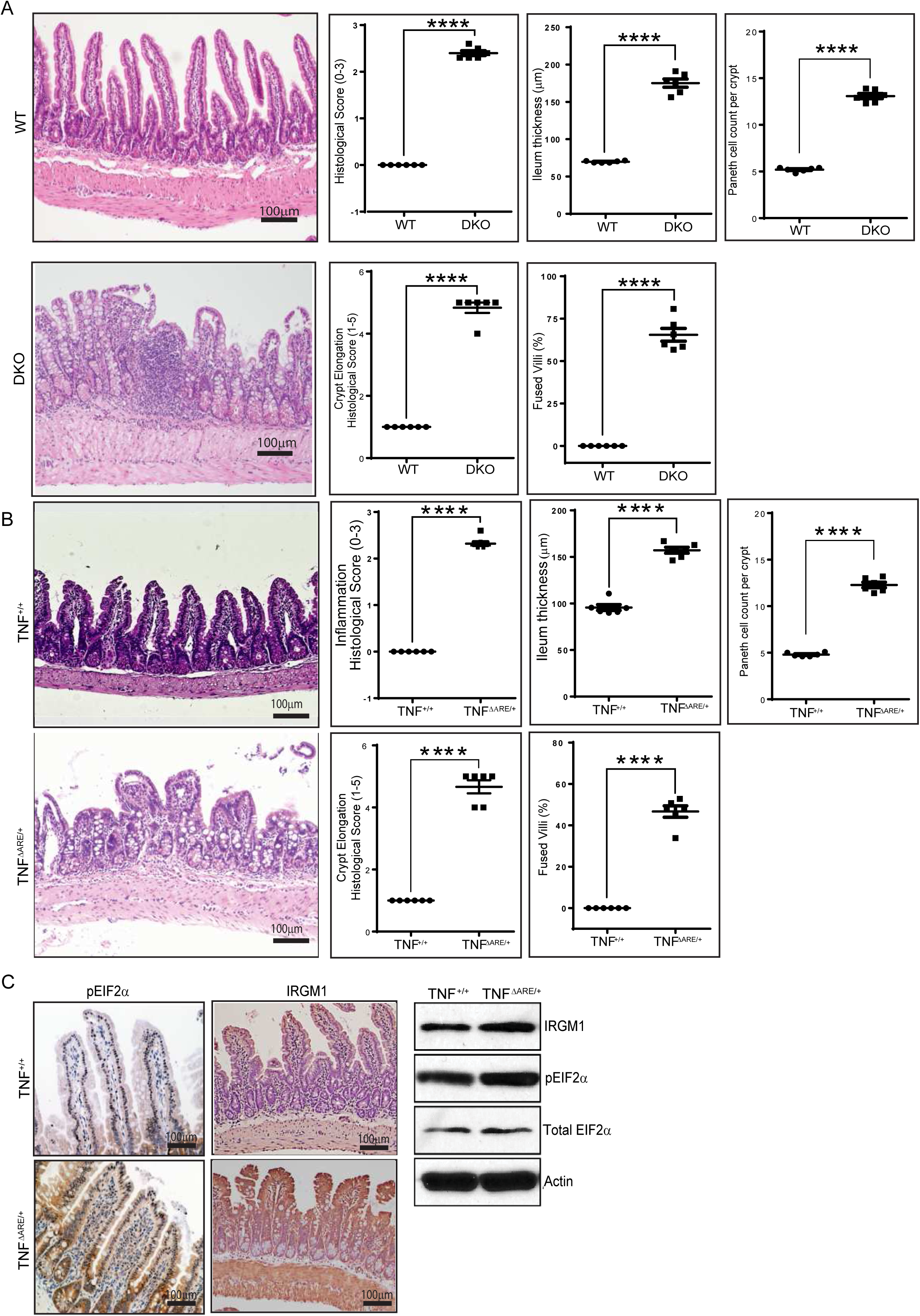
ISR activation is a hallmark of epithelial injury in murine models of CD. **(A-B)** Representative Hematoxylin and eosin (H&E) stained ileal sections from distal ileum of villin-1/gelsolin double knockout (DKO) and TNF^ΔARE/+^ mice and their wild-type (WT) littermates which includes TNF^+/+^ mice. DKO and TNF^ΔARE/+^ mice show ileal wall thickening, villus blunting, villus fusion, and crypt elongation compared to their respective control mice. Histology scores, measuring intestinal inflammation, were significantly elevated in DKO and TNF^ΔARE/+^ mice compared to their control mice. **(C)** Immunohistochemistry and immunoblots show upregulation of phospho-EIF2α (pEIF2α) and IRGM1 expression levels in the TNF^ΔARE/+^ mice without any notable change in total eIF2α levels, confirming chronic ISR activation. Data are presented as mean ± SEM from ≥3 independent experiments. Statistical significance was determined using two-tailed Student’s t-test, ****p<0.0001, n=6 mice per group.

At the molecular level, phosphorylated eIF2α (peIF2α), a marker of ISR activation, was strongly upregulated in TNF^ΔARE/+^ ileal tissue compared to TNF^+/+^ controls, whereas total eIF2α remained unchanged [Fig 1C]. IRGM1, a regulator of autophagy and mitochondrial homeostasis, was also markedly elevated in the TNF^ΔARE/+^ mice compared to the TNF^+/+^ mice. These results extend our prior findings in DKO mice and CD patients, demonstrating that despite distinct initiating stressors, epithelial intrinsic *versus* extrinsic and inflammatory, chronic ISR activation and IRGM1 induction are convergent hallmarks of epithelial injury in CD.[16]

### Chronic ISR impairs epithelial viability and regenerative capacity in murine and CD patient-derived enteroids

To directly assess epithelial cell-intrinsic responses, we derived enteroids from the distal ileum of DKO and TNF^ΔARE/+^ mice, as well as from ileal pinch biopsies from CD patients and non-CD controls. Phase contrast imaging and propidium iodide (PI) staining revealed significantly increased IEC death in DKO and TNF^ΔARE/+^ enteroids compared with their respective controls [Fig 2A-B]. Quantification showed markedly elevated PI-positive cells in DKO and TNF^ΔARE/+^ enteroids, confirming loss of membrane integrity. Necroptosis inhibition with Nec-1 rescued IEC survival in TNF^ΔARE/+^ mouse enteroids, whereas pan-caspase inhibition (Z-VAD-FMK) did not, confirming necroptosis rather than apoptosis as the dominant mode of cell death [Supplementary Fig 1] similar to our findings in the DKO mice and CD PDEs.[16] Enteroid forming efficiency, a measure of intestinal stem cell (ISC) regenerative potential, was profoundly reduced in both DKO and TNF^ΔARE/+^ mouse enteroids relative to control enteroids (p<0.0001, n=4 mice per group). Once formed enteroids from these murine CD models exhibited diminished complexity, with fewer crypt-like buds per enteroid and a reduced percentage of budded enteroids [Fig 2A-B]. These findings reflect defects in ISC-driven regeneration and epithelial renewal. Strikingly, CD patient-derived enteroids (PDEs) recapitulated the murine phenotypes, showing impaired survival, decreased enteroid forming efficiency, reduced bud number, and lower proportion of budded enteroids compared to non-CD controls [Fig 2C; p<0.0001, n=8 patients per group].

**Figure 2.**
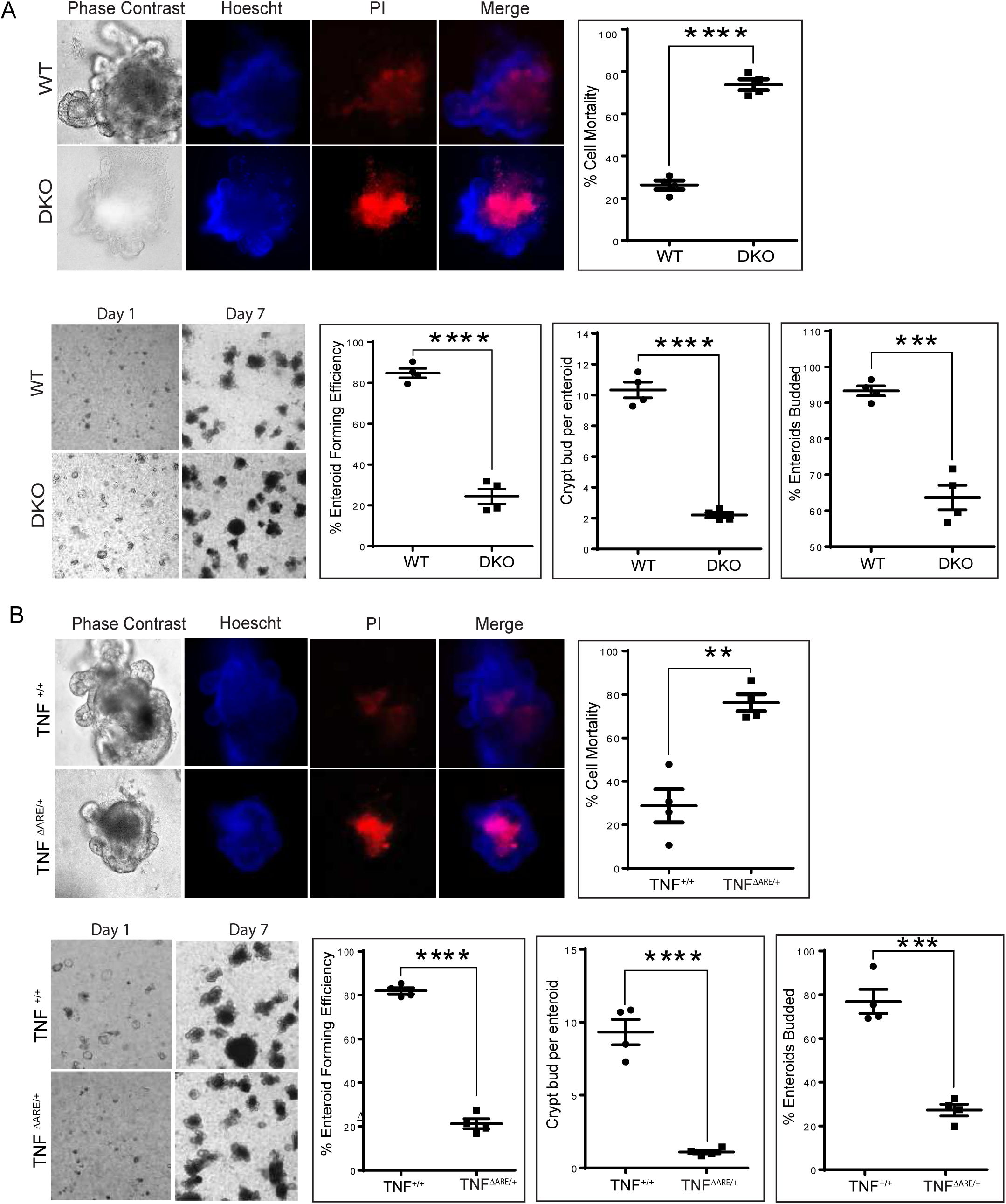
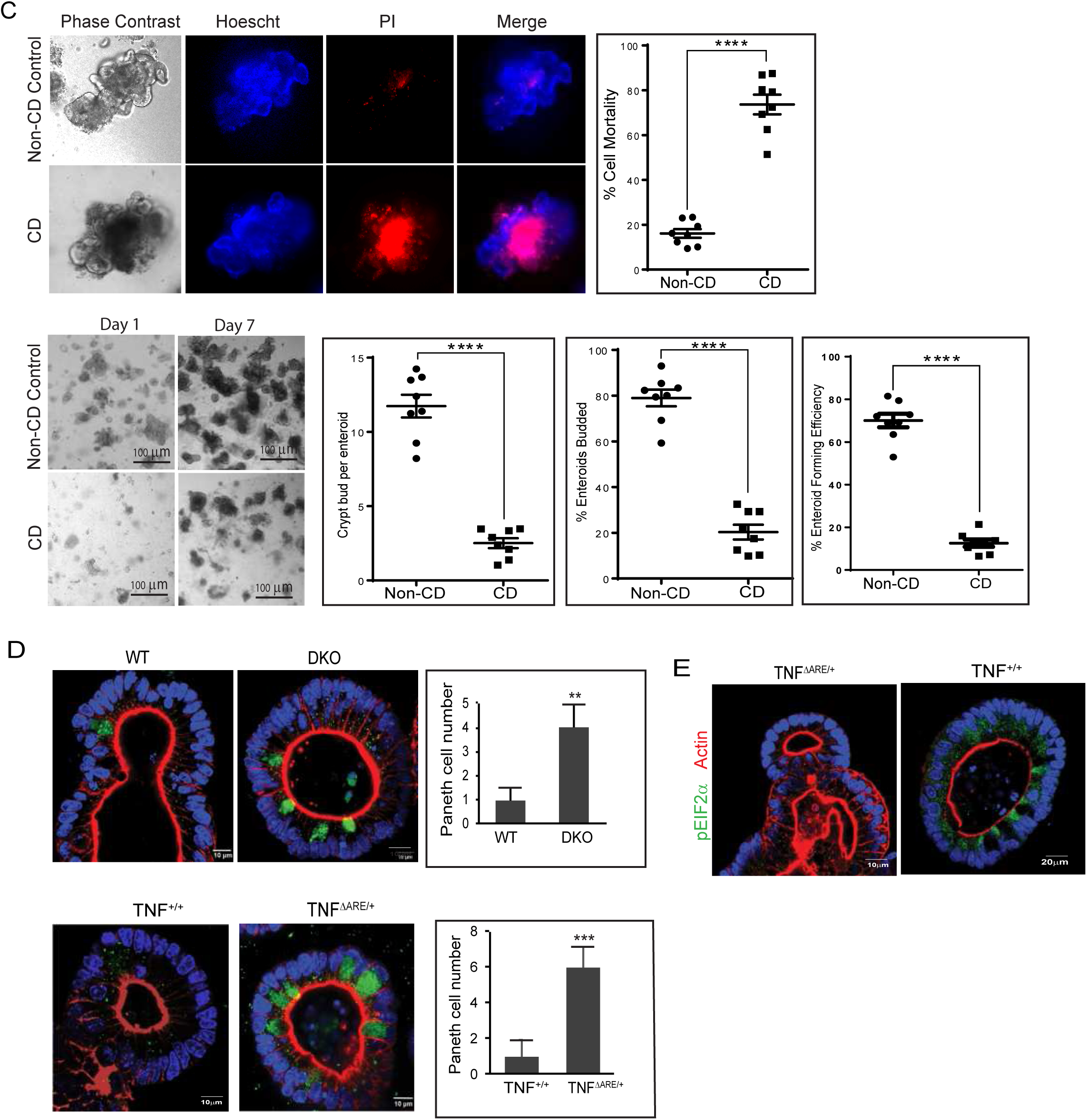
Chronic ISR activation impairs epithelial viability, regeneration, and growth in mouse and human models of CD. **(A-B)** Representative phase-contrast and fluorescence images of propidium iodide (PI) and Hoechst 33342 labelled enteroids from DKO, TNF^ΔARE/+^ and control mice (WT and TNF^+/+^, respectively) show increased intestinal epithelial cells (IEC) mortality in both murine models of CD. (A-B) Enteroids seeded at day 1 and analyzed at day 7 show significant inhibition of enteroid forming efficiency (percentage of plated crypts forming viable enteroids) in Controls versus DKO and TNF^ΔARE/+^ mice. Quantification of crypt buds per enteroid at Day 7 show reduced budding in DKO and TNF^ΔARE/+^ mouse enteroids compared to their respective controls. Percentage of budded enteroids is significantly decreased in DKO and TNF^ΔARE/+^ mice at Day 7 relative to controls. (**C)** CD patient-derived enteroids (PDEs) exhibit significantly increased mortality and inhibition of enteroid forming efficiency, number of enteroids budded, and total number of budded enteroids formed compared to PDEs from non-CD controls. **(D)** Representative confocal images show Paneth cell expansion in DKO and TNF^ΔARE/+^ mouse enteroids. Quantification of Paneth cell numbers reveals a significant increase in both DKO and TNF^ΔARE/+^ mouse enteroids compared to their respective control mouse enteroids. **(E)** Representative confocal images of enteroids immunoassayed for peIF2α shows significant upregulation of peIF2a expression in the TNF^ΔARE/+^ mouse enteroids compared to TNF^+/+^ mouse enteroids. Data obtained from 4 mice per group and 8 patients per group and expressed as mean ± SEM from ≥3 independent experiments. Statistical significance was assessed using one-way ANOVA with Tukey’s post-hoc test; **p<0.01, ***p<0.001, ****p < 0.0001. At least 50 enteroids per sample were analyzed for the quantitative analysis.

Confocal imaging revealed Paneth cell expansion in the DKO and TNF^ΔARE/+^ mouse enteroids, consistent with the *in vivo* phenotype [Fig 2D]. In addition, TNF^ΔARE/+^ enteroids exhibited sustained peIF2α expression compared to TNF^+/+^ control enteroids [Fig 2E]. Collectively, these results demonstrate that chronic ISR activation compromises epithelial survival, reduces ISC regenerative capacity, and drives Paneth cell lineage expansion across murine and human models of CD. Together with our prior work, these data establish necroptosis as a principal driver of IEC death and chronic ISR as a central regulator of regenerative failure in CD.[16]

### Pharmacological inhibition of ISR or necroptosis restores epithelial architecture in vivo

We next tested whether targeting ISR or necroptosis could mitigate epithelial pathology *in vivo*. DKO and TNF^ΔARE/+^ mice were treated with Nec-1 (RIPK1/3 inhibitor) or ISRIB, a small-molecule inhibitor of ISR that binds to eIF2β and restores protein synthesis even when eIF2α is phosphorylated.[35] Essentially, ISRIB “overrides” the ISR brake on translation.[36] Histological analysis of DKO mouse ileum revealed significant restoration of mucosal architecture in the Nec-1- and ISRIB-treated mice compared to vehicle-treated controls [Fig 3A-B, p<0.0001, n=6 mice per group.]. Drug-treated mice exhibited reduced villus fusion, normalized crypt length, and decreased muscularis propria thickening [p<0.0001, n=6 mice per group.]. Importantly, Paneth cell hyperplasia was significantly reduced following Nec-1 or ISRIB treatment of DKO mice, indicating correction of lineage imbalance. Histological scores confirmed marked suppression of intestinal inflammation [p<0.0001, n=6 mice per group]. We next asked whether ISR inhibition would provide similar benefit in an inflammation-driven model of Crohn’s-like ileitis. In TNF^ΔARE/+^ mice, treatment with ISRIB significantly reduced histological scores compared with vehicle controls [Fig 3C, p<0.0001, n=6 mice per group]. Additionally, ISRIB treatment led to significant reduction in villus fusion, crypt elongation, and ileal thickening, thus restoring the ileal tissue architecture [p<0.0001, n=6 mice per group]. This was accompanied by a significant reduction in the number of Paneth cell counts per crypt, mirroring the effects of ISRIB in DKO mice [p<0.0001, n=6 mice/group]. Together, these findings demonstrate that chronic ISR activation and necroptotic IEC death are tractable therapeutic vulnerabilities, and that pharmacologic inhibition of either pathway can restore epithelial homeostasis *in vivo*.

**Figure 3.**
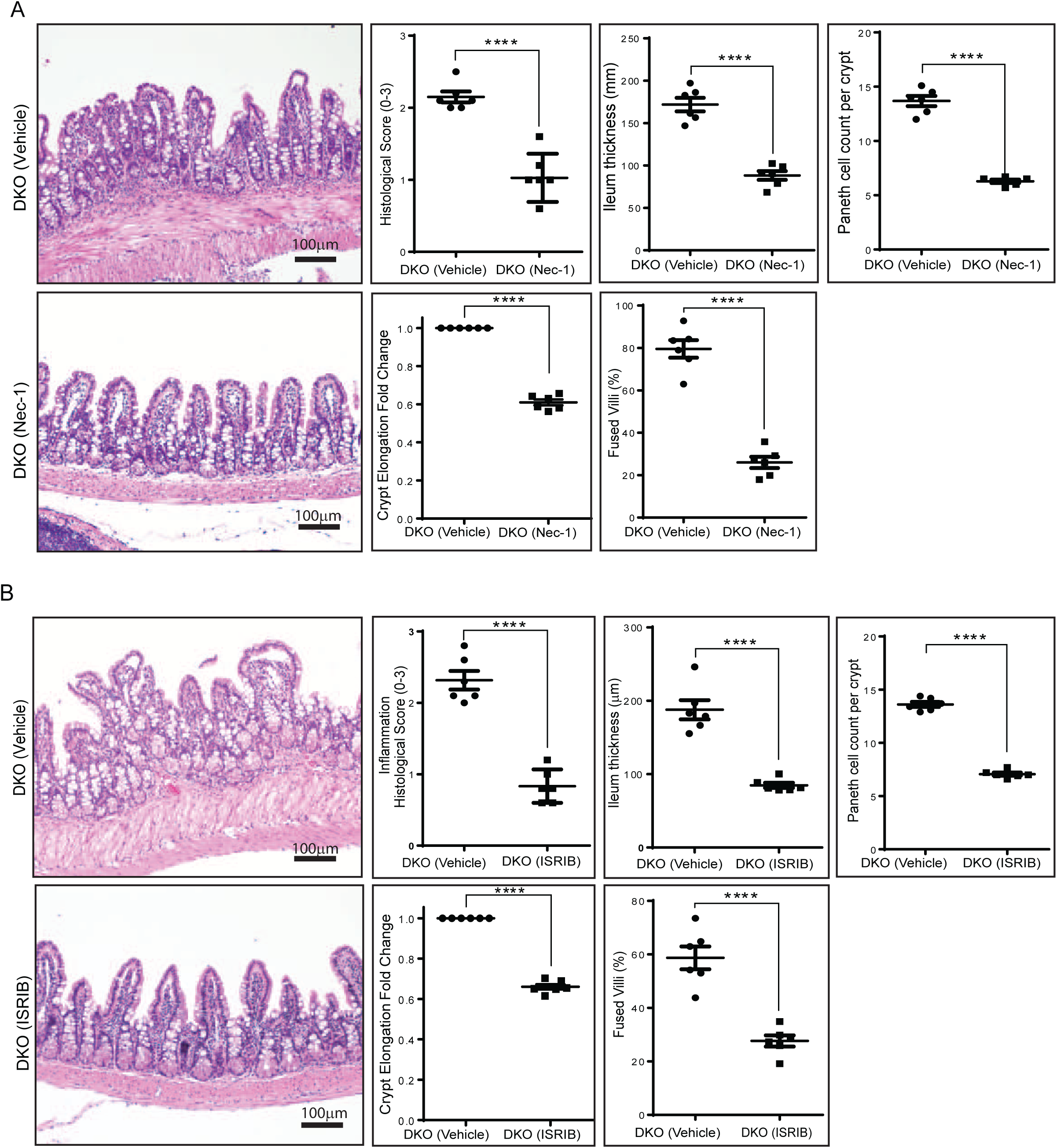
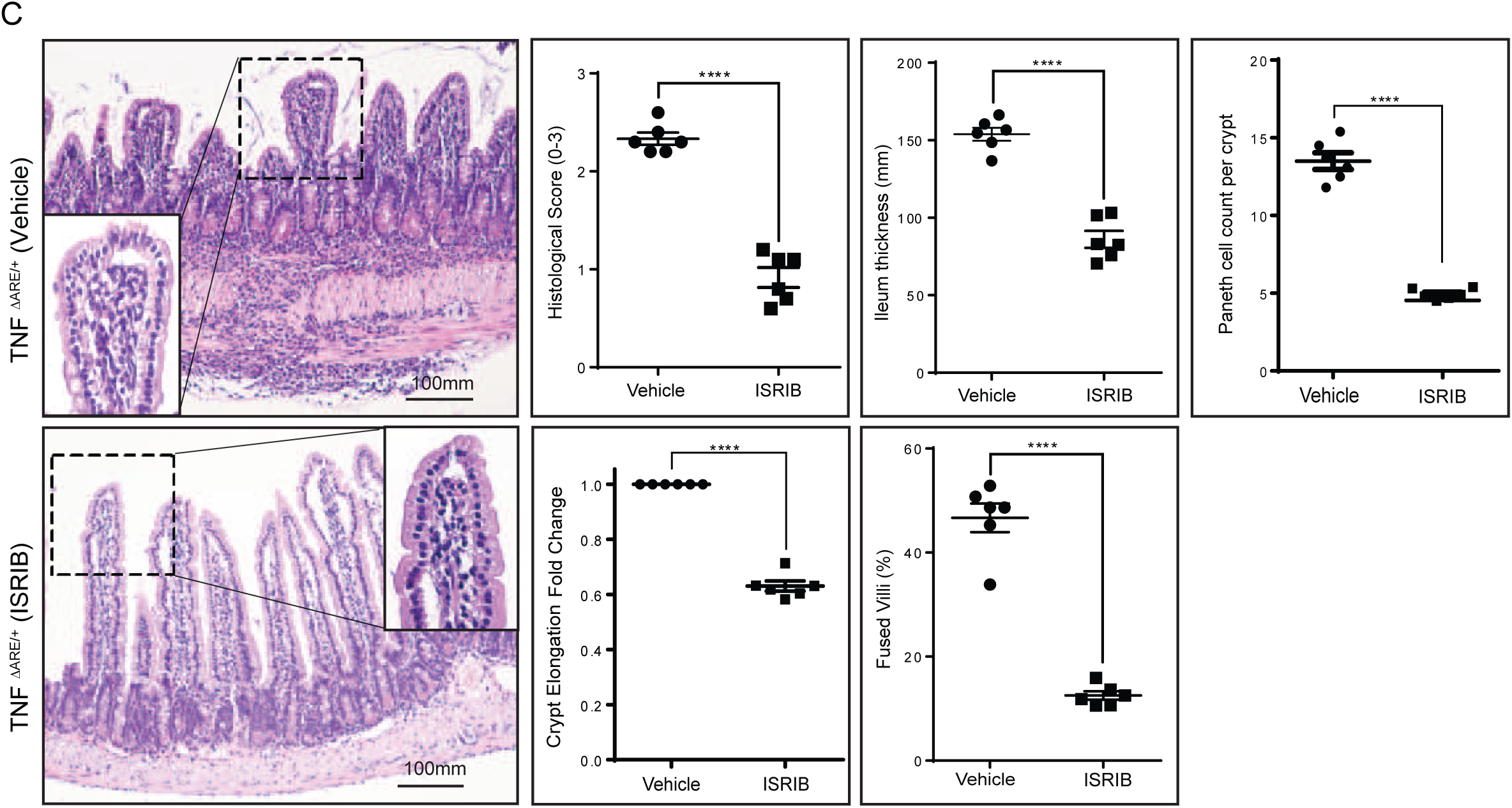
In vivo pharmacologic inhibition of ISR and necroptosis restore epithelial homeostasis in murine models of CD. DKO **(A-B)** and TNF^ΔARE/+^ **(C)** mice were administered ISRIB (0.25 mg/kg, i.p.) or Necrostatin-1 (Nec-1; 1.65 mg/kg; i.p.) as described in the Methods. Representative H&E-stained ileal sections show restoration of tissue architecture in the DKO, and TNF^ΔARE/+^ mice treated with Nec-1 and ISRIB compared to vehicle-treated controls. Both treatments significantly improved the histologic score showing inhibition of intestinal inflammation. Other indices of tissue histomorphology such as thickening of the muscularis propria, crypt elongation, number of fused villi and total numbers of Paneth cells per crypt, were also significantly reduced by Nec-1 and ISRIB treatment compared to vehicle-treated controls. Data is presented as mean ± SEM from ≥3 independent experiments. Statistical analysis was performed using one-way ANOVA with Tukey’s post-hoc test; ****p < 0.0001; n=6 mice per group.

### Repurposed cancer drugs with RIPK3 inhibitory properties rescue epithelial survival and regeneration in CD

Several FDA-approved cancer drugs including pazopanib, ponatinib, sorafenib, dabrafenib, ibrutinib, quizartinib, and regorafenib, have been found to selectively inhibit RIPK3.[37] Given the translational limitations of ISRIB and Nec-1, we tested FDA-approved cancer drugs pazopanib and ponatinib, both identified as potent RIPK3 inhibitors at sub-oncologic doses.[38] *Ex vivo* treatment of DKO and TNF^ΔARE/+^ mouse enteroids and CD PDEs with ISRIB (150 nM), Necrostatin-1 (10 μM), pazopanib (0.5 μM), or ponatinib (2 nM) markedly restored enteroid-forming efficiency, crypt budding, and increased the number of budded enteroids, while also reducing IEC mortality compared with their respective vehicle (DMSO)-treated controls [Fig 4A-C; p<0.0001, enteroids were derived from n=4 mice and n=6 patients; Supplementary Fig 2.]. The magnitude of rescue with pazopanib and ponatinib was comparable to that observed with ISRIB and Nec-1, indicating comparable efficacy. These results show that pazopanib and ponatinib can rescue epithelial viability and regenerative capacity in both murine and human CD enteroids, highlighting their potential as clinically translatable mucosal healing therapies in CD.

**Figure 4.**
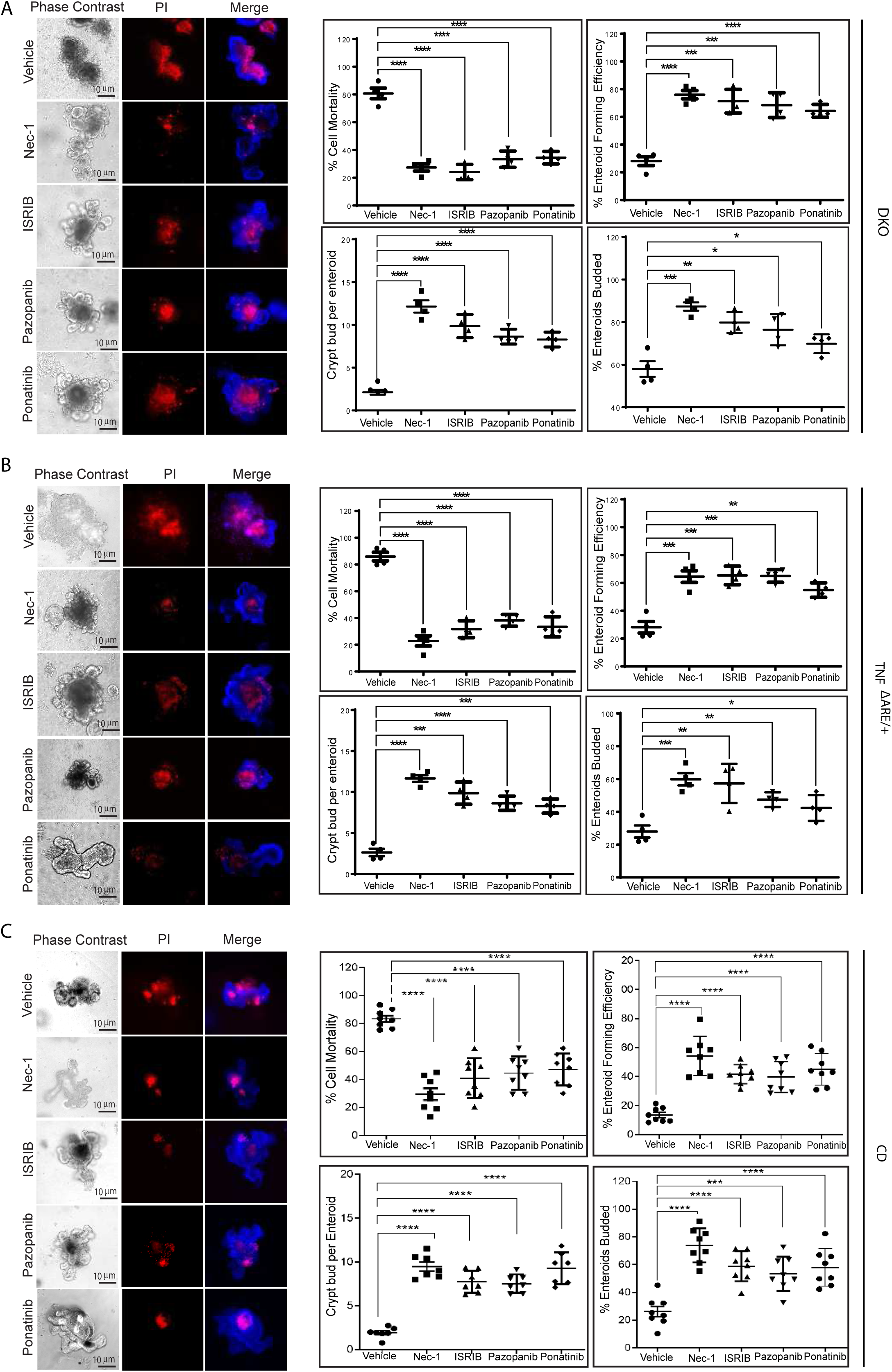
Repurposed cancer drugs with potent RIPK3 inhibiting properties, rescue epithelial viability and regeneration. **(A-C)** Representative phase-contrast and PI stained-merged images of DKO and TNF^ΔARE/+^ and CD PDEs treated with ISRIB, Nec-1, pazopanib, ponatinib, or vehicle (DMSO) show significant inhibition of IEC mortality. Enteroids exogenously treated with these inhibitors also significantly improved enteroid forming efficiency, number of enteroids with buds, and the number of buds per enteroid compared to vehicle-treated control enteroids. Data is from 4 mice per group and 8 patients per group, represented as mean ± SEM from ≥3 independent experiments. Statistical significance was assessed using one-way ANOVA with Tukey’s post-hoc test; asterisks (*) represent: **p<0.01, ***p<0.001, ****p< 0.0001.

### Repurposed RIPK3 inhibitors restore permeability barrier integrity in CD PDE

Finally, we evaluated whether these pharmacological interventions improved functional permeability barrier integrity, as measured by transepithelial electrical resistance (TEER). Treatment with pazopanib, ponatinib, ISRIB, or Nec-1 significantly improved TEER values compared to vehicle-treated controls [Fig 5, p<0.001, n=6 patients per group.]. In CD PDE monolayers, pazopanib and ponatinib significantly improved transepithelial electrical resistance (TEER), directly demonstrating functional recovery of permeability barrier integrity. These results directly demonstrate that repurposed RIPK3 inhibitors not only promote mucosal healing but also promote permeability barrier healing in human CD epithelium.

**Fig. 5.**
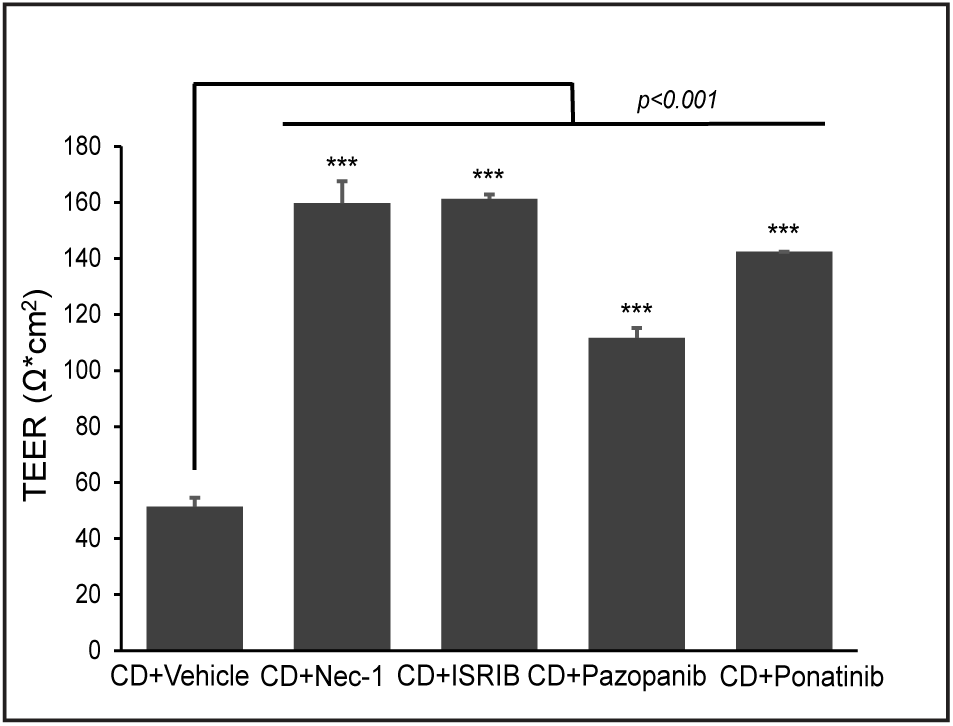
Pharmacologic blockade of ISR, necroptosis, and RIPK3 signaling restore epithelial permeability barrier function in CD patient-derived enteroids. Transepithelial electrical resistance (TEER, Ω.cm^2^) was measured in CD PDE treated with vehicle (DMSO), Nec-1, ISRIB, pazopanib, or ponatinib. All agents significantly improved TEER compared with vehicle-treated CD enteroids, indicating restoration of epithelial barrier integrity. Data represent mean ± SEM from ≥3 independent experiments; asterisks (*) denote: ***p<0.001.

Across murine models and human CD PDEs, we show that: (1) chronic ISR activation and necroptosis are convergent hallmarks of epithelial dysfunction; (2) these pathways drive IEC death, regenerative failure, IEC lineage defects, and permeability barrier loss; (3) pharmacologic inhibition with ISRIB, Nec-1, pazopanib, or ponatinib rescues epithelial survival, regenerative growth, and permeability barrier integrity. These findings position ISR and RIPK3 inhibition as promising therapeutic strategies to achieve permeability barrier healing in CD.

## Discussion

Our results highlight ISR activation as a unifying hallmark of epithelial stress, regardless of whether the initiating insult is intrinsic (loss of villin-1/gelsolin) or extrinsic (TNF-α dysregulation). TNF^ΔARE/+^ mice display chronic peIF2α phosphorylation and IRGM1 induction, molecular features we previously identified in the DKO mice and CD patients.[16] By regulating mitochondrial fission and membrane polarization, IRGM1 influences mitochondrial quality control while overexpression of IRGM1 causes mitochondrial depolarization and necroptotic cell death as we have shown in DKO mice and CD patients.[16, 39] In acute stress, the ISR restores homeostasis and promotes survival.[15] However, our data demonstrate that chronic ISR activation in the gastrointestinal tract is maladaptive, locking epithelial cells in a state of translational arrest and mitochondrial stress that culminates in necroptotic cell death and impairment of epithelial regeneration.[16] By showing that ISR activation and RIPK3-regulated necroptosis converge across distinct etiologies, we identify a common pathogenic axis that explains persistent epithelial dysfunction in CD and offers a tractable therapeutic target.

In this study, enteroid assays provided critical functional insights into how ISR and necroptosis compromise epithelial homeostasis and function. The parallel defects in enteroid formation, budding, and regenerative capacity across murine and CD PDEs underscores the role of chronic ISR activation in compromising ISC survival and epithelial regenerative potential. In response to injury, stem cells adopt new fates and lineages to support regeneration of the damaged tissue. In the epidermis, ISR activation determines whether skin stem cells maintain their identity or undergo a lineage conversion thus playing a key role in the re-epithelialization of the injured epidermis.[40] However, in aging or chronic stress conditions, the ISR contributes to dysfunctional epithelial regeneration, as seen in the lung tissue where ISR hinders effective repair resulting in fibrosis.[41] Our study shows chronic ISR activation in CD impairs ISC renewal and epithelial regeneration thus directly contributing to defects in mucosal healing and permeability barrier dysfunction in CD. Additionally, together with our prior work, in this study we show aberrant Paneth cell expansion in CD.[16] Paneth cells provide essential niche signals for stem cell maintenance, and antimicrobial peptides and their expansion may represent a compensatory but dysfunctional response to epithelial stress. These findings reinforce the concept that chronic ISR disrupts epithelial homeostasis at multiple levels, by depleting viable IECs but also by inhibiting ISC renewal, disrupting lineage specification, and crypt-villus dynamics. ISR activation has been linked to cellular metabolism and in the epidermis, ISR modulates stem cell fate decisions in response to serine metabolism.[40] This raises the intriguing possibility that dietary interventions could synergize with ISR-targeted therapies to promote mucosal and permeability barrier healing in CD. Together with our previous findings, data provided here show that in CD, pathologic IEC death in the intestine is mediated primarily by necroptosis and not apoptosis.[16] In the small intestine, enterocytes and enterocyte progenitor cells are particularly sensitive to necroptosis, a caspase-independent lytic form of programmed cell death.[16, 42, 43] We postulate that the increased susceptibility of the enterocyte progenitors to necroptosis, contributes to impaired epithelial regeneration identified in the DKO, TNF^ΔARE/+^ and CD PDEs.

The link between impaired gut barrier and inflammatory bowel disease (IBD) was established >30 years ago, but the lack of clinical translation is in part due to the lack of identifiable therapeutic targets. Previous studies have shown that growth factors such as EGFR and R-spondin1 can restore the damaged intestinal epithelium, however both are mitogens and could increase the risk of developing colorectal cancer.[44, 45] Larazotide acetate was identified as a BH therapy however, its barrier-protective effects were not recapitulated in clinical trials.[46] A major advance of this work is the demonstration that pharmacologic inhibition of ISR or necroptosis restores epithelial homeostasis by promoting regenerative growth of the epithelium in both murine and human models of CD. *In vivo*, both ISRIB and Necrostatin-1 significantly improved villus architecture, reduced inflammation, and normalized Paneth cell numbers in DKO and TNF^ΔARE/+^ mice. *Ex vivo*, ISRIB and Nec-1 improved IEC viability, ISC renewal, and regeneration of the crypt-villus-like enteroid architecture. Notably, ISRIB promotes lung repair and reduces fibrosis by accelerating the differentiation of alveolar epithelial cells after injury.[47] This implies the potential of ISRIB to restore epithelial homeostasis across different tissues. It also indicates that given prophylactically, ISRIB could reduce or prevent fibrostenotic strictures in CD. Treatment of structuring and fistulizing CD is one of the hardest challenges in CD care.[48] Strictures are caused by chronic inflammation and no approved antifibrotic drug exists yet. Moreover, once fibrosis is established in the gut, treatment has limited impact. While anti-TNF biologics can reduce inflammatory components of a stricture, they do not reverse scar tissue and surgical management remains the only option. Notably, the ERICa trial demonstrated that in CD patients, permeability barrier healing is associated with a decreased risk of developing intestinal complications.[2] Consistent with that, we propose that inhibition of ISR has the potential to prevent the initiation and development of fibrostricturing CD. ISRIB, though investigational, has shown safety and oral bioavailability in preclinical studies, making it attractive for translational development for CD.[49] Pazopanib and ponatinib are FDA-approved with well-characterized pharmaco-kinetics.[38] At significantly low doses, both drugs selectively inhibit RIPK3 without causing systemic toxicity.[38, 50] We show that at sub-oncologic doses, pazopanib and ponatinib effectively restore epithelial viability, epithelial regenerative capacity, and TEER in CD PDEs. Unlike investigational RIPK3 inhibitors like GSK’872 and GSK843, which can paradoxically trigger apoptosis, pazopanib and ponatinib have been shown to suppress both necroptosis and RIPK3-driven inflammatory signaling, making them ideal candidates for CD therapy.[51] We postulate that the dual effect of pazopanib and ponatinib on necroptosis and inflammation may be critical for effective restoration of permeability barrier integrity in CD.

ER in CD is associated with decreased risk of disease progression and a more favorable long-term disease outcome.[10] Multiple studies have used confocal endomicroscopy to assess the structure and function of the intestinal permeability barrier *in vivo* and demonstrated that impaired barrier integrity in CD patients is linked with persistence of clinical symptoms and relapsing disease behavior.[2, 13, 14, 52] These findings have led to the idea that mucosal healing or endoscopic remission is not sufficient to improve treatment outcomes and that restoration of permeability barrier is essential to achieve sustained clinical remission and to reduce intestinal complications in CD. Despite that, there are no clinically validated methods for early diagnosis, risk stratification, or targeted intervention based on impaired permeability and there are no BH therapies. Our study introduces a paradigm shift in CD therapy, by focusing on BH, an unaddressed yet decisive driver of clinical outcomes in CD.

Our work also opens several avenues for further investigation. First, development of gut-restricted RIPK3 inhibitors could maximize efficacy while minimizing systemic exposure. Second, incorporating BH as a therapeutic endpoint in clinical trials will be critical for aligning CD therapy with outcomes most predictive of long-term health.[2] Additionally, biomarkers of ISR activation and RIPK3 signaling, such as peIF2α or RIPK3 expression in mucosal biopsies, could be used to stratify patients and guide therapy. Identifying patients with persistent barrier dysfunction will allow targeted use of barrier-restoring agents. Finally, integration of PDE-based assays and TEER measurements into personalized medicine pipelines could enable *ex vivo* prediction of treatment response in CD.

### Conclusions

Our study defines ISR activation and RIPK3-mediated necroptosis as convergent mechanisms driving epithelial dysfunction in CD. By demonstrating that pharmacological inhibition of these pathways restores epithelial regeneration and permeability barrier integrity, we provide proof-of-concept for targeting epithelial stress response and necroptosis as a therapeutic strategy. Repurposing pazopanib and ponatinib offers an immediately translatable approach to achieve permeability barrier healing, a treatment endpoint superior to endoscopic healing for predicting long-term disease outcomes. Incorporating permeability barrier healing into therapeutic targets and clinical trial design has the potential to fundamentally reshape CD management, moving beyond inflammation control toward sustained restoration of epithelial homeostasis.

---

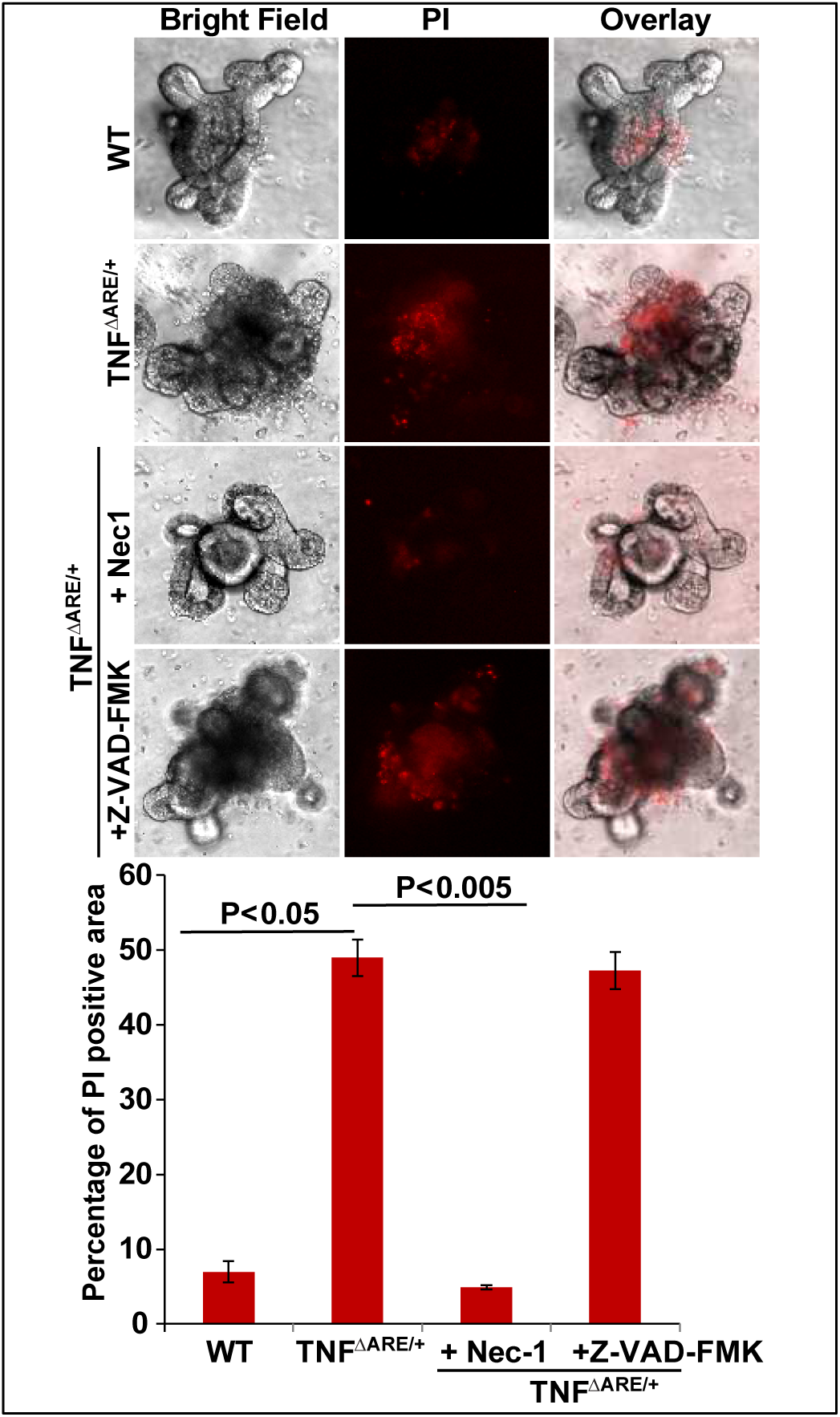

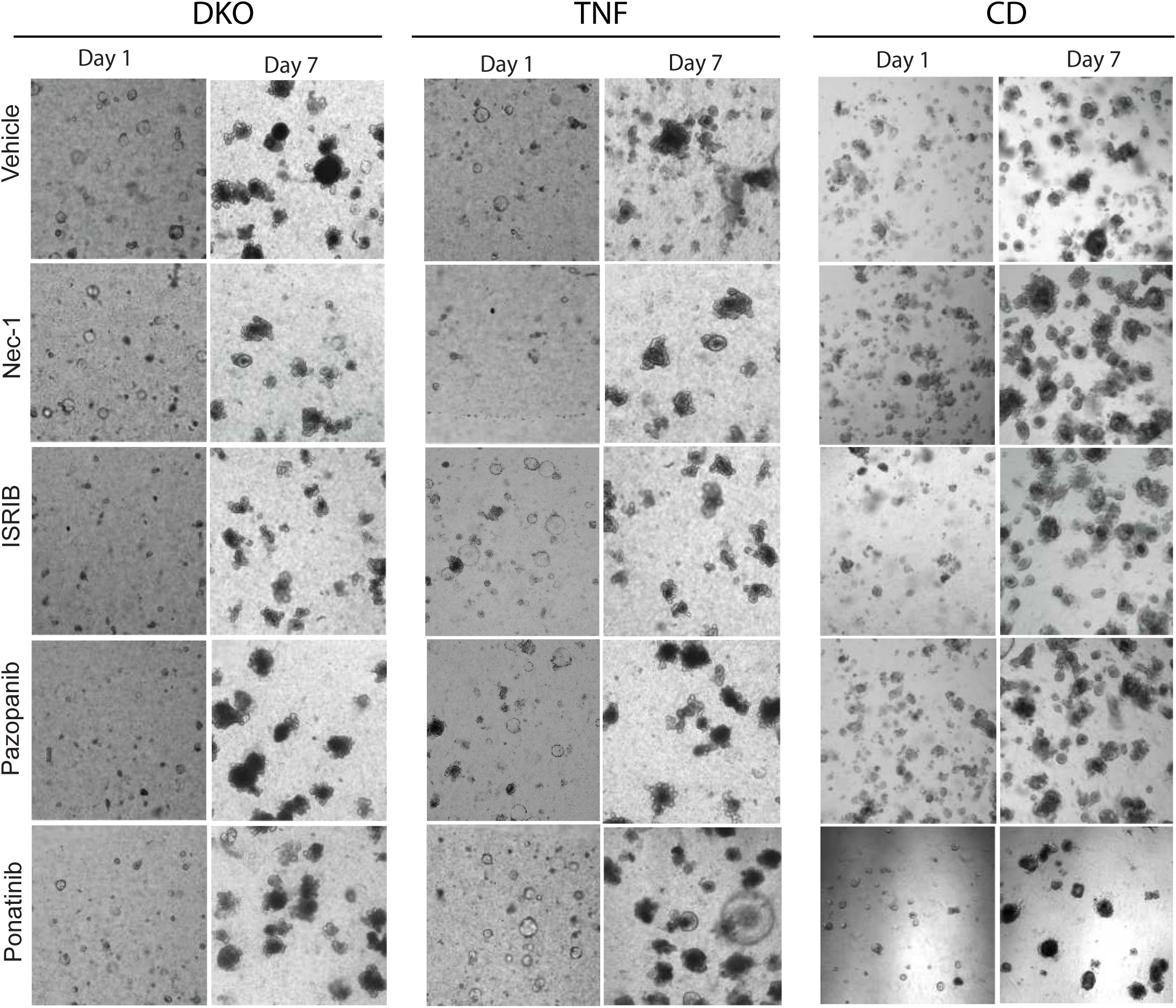

## Notes

**<u>Grant</u>**<u>**Support**</u>: Supported by NIDDK (DK-98120, DK-117476 to S.K.) and the Public Health Service (DK-056338).

### Competing Interest Statement

The authors have declared no competing interest.

